# Astrocytic glutamate uptake coordinates experience-dependent, eye-specific refinement in developing visual cortex

**DOI:** 10.1101/2020.05.25.113613

**Authors:** Grayson Sipe, Jeremy Petravicz, Rajeev Rikhye, Rodrigo Garcia, Nikolaos Mellios, Mriganka Sur

## Abstract

The uptake of glutamate by astrocytes actively shapes synaptic transmission, however its role in the development and plasticity of neuronal circuits remains poorly understood. The astrocytic glutamate transporter, GLT1 is the predominant source of glutamate clearance in the adult mouse cortex. Here, we examined the structural and functional development of the visual cortex in GLT1 heterozygous (HET) mice using two-photon microscopy, immunohistochemistry and slice electrophysiology. We find that though eye-specific thalamic axonal segregation is intact, binocular refinement in the primary visual cortex is disrupted. Eye-specific responses to visual stimuli in GLT1 HET mice show altered binocular matching, with abnormally high responses to ipsilateral compared to contralateral eye stimulation and a greater mismatch between preferred orientation selectivity of ipsilateral and contralateral eye responses. Furthermore, the balance of excitation and inhibition in cortical circuits is dysregulated with an increase in somatostatin positive interneurons, decrease in parvalbumin positive interneurons, and increase in dendritic spine density in the basal dendrites of layer 2/3 excitatory neurons. Monocular deprivation induces atypical ocular dominance plasticity in GLT1 HET mice, with an unusual depression of ipsilateral open eye responses; however, this change in ipsilateral responses correlates well with an upregulation of GLT1 protein following monocular deprivation. These results demonstrate that a key function of astrocytic GLT1 function during development is the experience-dependent refinement of ipsilateral eye inputs relative to contralateral eye inputs in visual cortex.

**SIGNIFICANCE:** We show that astrocytic glutamate uptake via the transporter GLT1 is necessary for activity-dependent regulation of cortical inputs. Dysregulation of GLT1 expression and function leads to a disruption of binocular refinement and matching in visual cortex. Inputs from the ipsilateral eye are stronger, and monocular deprivation, which upregulates GLT1 expression in a homeostatic fashion, causes a paradoxical reduction of ipsilateral, non-deprived eye, responses. These results provide new evidence for the importance of glutamate transport in cortical development, function, and plasticity.

## INTRODUCTION

Astrocytes constitute a major class of cells in the mammalian brain with critical roles in brain homeostasis, function, and plasticity (Verkhratsky and Nedergaard, 2018). However, the role of astrocytes in cortical development and plasticity remains poorly understood. Astrocyte processes are known to form intimate contacts with neuronal synapses ensheathing sites of neurotransmitter release and regulating synaptic efficacy and plasticity (Eroglu and Barres, 2010). Furthermore, astrocytes are the primary cell type responsible for clearance of synaptic glutamate via excitatory amino acid transporters (EAATs) (Danbolt, 2001). Neuronal excitability and synaptic transmission are modulated by the expression of EAATs in numerous brain regions including the cortex (Campbell and Hablitz, 2004), thalamus (Hauser et al., 2013), spinal cord (Weng et al., 2007), cerebellum (Takatsuru et al., 2007) and hippocampus (Huang et al., 2004). Astrocytes express two EAATs, GLAST (EAAT1) and GLT1 (EAAT2) with varying regional expression during development to adulthood (Danbolt, 2001). In the adult mouse cortex, GLT1 is responsible for approximately 80-90% of synaptic glutamate clearance, thereby representing a critical mechanism for modulating synaptic transmission and function (Danbolt, 2001). In addition, the dynamic ensheathment of synapses by astrocyte processes (Perez-Alvarez et al., 2014), the modulation of GLT1 expression by increased network activity (Murphy-Royal et al., 2015), and the subcellular trafficking of GLT1 to highly active synaptic zones (Benediktsson et al., 2012) together suggest that GLT1 serves a critical role in astrocyte-neuron signaling so as to facilitate synaptic plasticity through dynamic glutamate uptake commensurate with synaptic strength. As such, disruptions in GLT1 expression or function have been implicated in several neuropathologies (Cui et al., 2014; Sugimoto et al., 2018; Robinson et al., 2020). However, the role of astrocytic GLT1 in early brain development and functional maturation remains inadequately characterized.

The mouse primary visual cortex (V1) has served as a useful model system for studying cortical development and plasticity (Hensch, 2005; Schummers et al., 2005; Leamey et al., 2009; Hooks and Chen, 2020). Layer 2/3 excitatory neurons in the binocular region of V1 receive information from both the contralateral and ipsilateral eye via thalamocortical and intracortical inputs that are shaped during critical periods of experience-dependent plasticity (Hooks and Chen, 2020). The matching of binocular orientation preference, in which neurons receiving binocular input alter their synaptic connections to align the orientation preference of both contralateral and ipsilateral eye responses, represents one key example of experience-dependent plasticity (Antonini et al., 1999; Kara and Boyd, 2009; Wang et al., 2010; Bhaumik and Shah, 2014; Gu and Cang, 2016; Tie et al., 2018). This refinement of binocular matching is dependent upon NMDA receptors acting as coincidence detectors of contra- and ipsilateral drive (Sawtell et al., 2003), which are known to be modulated by GLT1 (Aida et al., 2012; Aida et al., 2015; Pinky et al., 2018). In addition, the anatomical and synaptic inputs onto binocular V1 neurons are weighted such that there is normally a contralateral eye bias in response amplitude (Antonini et al., 1999). Monocular deprivation (MD) of the contralateral eye for a few days (between ~P21 and 30) leads to decreased contralateral eye responses followed by a homeostatic increase in ipsilateral eye responses (Hensch, 2005; Hooks and Chen, 2020). This process is termed ocular dominance plasticity (ODP) and defines a second type of experience dependent plasticity in V1 (Katz and Crowley, 2002; Espinosa and Stryker, 2012; Trachtenberg, 2015).

In order to explore astrocytic roles in development via GLT1-mediated glutamate uptake, we characterized the developmental timeline of GLT1 expression and used a transgenic mouse line with constitutive heterozygous expression of the GLT1 gene (SLC1A2, GLT1-HET (Kiryk et al., 2008)) to examine the consequences of decreased astrocyte glutamate uptake in visual cortex development. Our data show that GLT1 heterozygosity results in disrupted binocular matching, abnormal excitation/inhibition, and aberrant ODP. Interestingly, large-scale synaptic remodeling induced by monocular deprivation causes a haplosufficient increase in GLT1 expression and blocks ipsilateral eye response potentiation, indicating that the ipsilateral inputs are particularly sensitive to astrocyte-mediated glutamate uptake. This work demonstrates an important and unexpected role for astrocytic glutamate uptake in the development and experience-dependent plasticity of eye-specific responses in V1.

## METHODS

### Animals and Surgery

GLT1-HET mice on a C57/Bl6 background were obtained from Kohichi Tanaka and Jeffrey Rothstein, and maintained in their home cage until experiments under a 12/12 Light/Dark cycle with food and standard mouse chow provided *ad libitum*. GLT1-HET mice were also crossed with Thy-GFP mice (Jackson labs) and GFAP-tDTomato mice to analyze neuronal and astrocyte morphology respectively. Male and female mice were used throughout the study with GLT1-WT littermates as controls. For in vivo imaging of neurons, GLT-1 Het mice were crossed to Emx-Cre::GCaMP6f mice (Jackson labs) to allow for imaging of neuronal responses in the binocular region of the visual cortex at P30-32. At age P24-25, mice were anesthetized using isoflurane (3% induction, 1.5–2% during surgery). A 3-mm-diameter craniotomy was performed over binocular V1 (2–3 mm lateral and 0.5 mm anterior to lambda). The craniotomy was covered with a 3 mm glass coverslip (Warner Instruments), and a custom-built metal head post was attached to the skull and sealed with dental cement (C&B-Metabond, Parkell). Care was taken not to rupture the dura mater. The core body temperature was maintained at 37.5°C using a heating blanket (Harvard Apparatus). For optical dominance plasticity experiments, MD was performed by eyelid suture. Animals (~P21-24) were anesthetized with isoflurane (3% induction, 1.5–2% during surgery), and eyelid margins trimmed. Upper and lower lids were sutured closed, and eyelids were regularly examined to ensure that they remained closed for the duration of the experiment. MD lasted either 4 or 7 days according to established models of short and long-term ODP (Nagakura et al., 2014; Hooks and Chen, 2020). Before optical imaging, the sutures were removed and the deprived eye reopened while the animal was under anesthesia. All animal procedures were performed in strict accordance with protocols established with the Massachusetts Institute of Technology, Division of Comparative Medicine, and conformed to NIH guidelines.

### Immunohistochemistry

Mice were deeply anesthetized using isoflurane and transcardially perfused with 0.1M phosphate buffered saline (PBS) and 4% paraformaldehyde in PBS (PFA). Whole brains were removed and post-fixed overnight at 4°C until sectioning. Coronal brain slices were acquired at a thickness of 40μm using a vibratome (VT1200S; Leica) and stained or directly slide-mounted with general purpose mounting media with DAPI (Vector Labs). Free-floating brain slices were washed (3×10min @ ~20°C, 0.1M PBS) and then blocked (1h at ~20°C; 5% BSA /1% triton in wash buffer). Immediately following blocking, tissue was incubated in primary antibody solution (24hr at ~4°C; 3% BSA/ 0.1% triton in wash buffer), washed (3×10min) and placed in secondary antibody solution (4hr at ~20°C, 3% BSA/ 0.1% triton in wash buffer). Tissue was then washed (3×10min) and mounted on to microscope slides as described above. The following primary antibodies and dilutions were used: rabbit α-GLT1 (1:500, AGC-022, Alamone Labs), guinea pig α-GFAP (1:2000, 173 004, Synaptic Systems), chicken α-GFP (1:1000, GFP-1020, Aves Labs), α-rabbit α-SST-14 (1:500, T-4103, Peninsula Labs), mouse α-PV (1:500, MAB1572, EMD Millipore). The following secondary antibodies and dilutions were used (1:500, AlexaFluor, Invitrogen): donkey α-mouse, goat α-rabbit, goat α-chicken and goat anti-guinea pig. For perineuronal net imaging, sections were stained using *wisteria floribunda agglutinin* (WFA), which labels chondroitin sulfate proteoglycan chains that constitute perineuronal nets, using a standard streptavidin/biotin kit (Vector Labs, SP-2002).

### mRNA and Protein Quantification

For quantification of GLT1 mRNA, mouse RNA from V1 was extracted using the miRNeasy Mini Kit (Qiagen) based on manufacturer’s instructions. RNA concentration and quality was assayed using NanoDrop (Thermo Scientific) and 300 ng of RNA were reverse transcribed using the Vilo cDNA kit (Invitrogen) with a 1:15 dilution of cDNA. GLT1 mRNA was normalized to β-actin using the following formula: GLT1 relative expression = 2^CT^β-actin^/2^CT^GLT1^ mRNA. For protein measurements of GLT1 and GLAST, V1 contralateral to the deprived eye was removed and snap-frozen with dry ice. The brains were homogenized in RIPA buffer (Invitrogen) containing proteinase and phosphatase inhibitors (Roche), centrifuged for 10 min at 8,000 × g at 4°C and supernatant extracted and stored at −80°C. Protein concentration was assayed using BCA protein assays (Pierce, Thermo Scientific). For Western blot, 10μg/μl of protein was loaded into each lane of 4–15% (mg/100 mL) Tris·HCl polyacryamide gels (Bio-Rad). Protein was transferred to Immobilon-P PVDF membranes (Millipore), blocked with 5% (mg/100 mL) BSA (Sigma) for 1 h, and incubated in the following antibody solutions overnight: GLT1 (1:25k, AB1783, EMD Millipore), GLAST (1:10k, ab181036, Abcam) and β-actin (1:20k, A1978; Sigma-Aldrich). Blots were then incubated in the following horseradish peroxidase-conjugated secondary antibodies (IRDye® 800CW Donkey anti-Rabbit IgG, Licor System) for 1.5 hr, washed and then imaged. Optical densities of detected bands were quantified using ImageJ software. A standard sample of wild-type mouse V1 tissue was run on each gel to gauge blot-to-blot variability.

### Labeling of Retinal Ganglion Axons

GLT1-HET and WT mice (~P28-P32) were anesthetized with isoflurane, and 2 μl of cholera toxin subunit B (CTB) conjugated to Alexa Fluor 488 was injected into the ipsilateral eye and Alexa Fluor 594 into the contralateral eye (1 mg/ml; Invitrogen). After 6 d, animals were perfused and prepared as described above. Projection overlap was quantified in ImageJ as previously described (Stevens et al., 2007; Nagakura et al., 2014; Ip et al., 2018).

### Two-photon Imaging

Two-photon *in vivo* imaging was performed as described previously (Rikhye and Sur, 2015). Images of GCaMP6f-positive neurons were obtained using a Prairie Ultima two-photon system (Bruker) driven by a Spectra Physics Mai-Tai laser passed through a Deep-See module (Spectra Physics) and a high-performance objective lens (25 Olympus XL Plan Nobjective, 1.05 numerical aperture). Cells were excited 910 nm for GCaMP6f. A custom-built MATLAB-based (MathWorks) software system was used to collect optimized raster scans at 50 frames/s and to perform offline data analysis. The binocular region of V1 was determined by covering the eye contralateral to the craniotomy with an eye patch, and neuronal responses elicited by displaying moving gratings. If neuronal responses were visually detected, responses were confirmed using the contralateral eye while the ipsilateral eye was covered. Fields where neuronal responses were detected to both contra- and ipsilateral stimulation were used in experiments. Single eye responses to orientated gratings were then collected from each field. To assess the orientation selectivity and tuning of neurons, we presented oriented gratings on a 23” 1080p LCD monitor (Dell) using custom software (Network Visstim, Sur Lab) written in PsychToolbox-3 (Psychtoolbox.com) on a Windows 10 computer (Dell Precision) with a GeForce 8800 GTS 640MB graphics card (PNY). Gratings were optimized for cellular responsiveness using a contrast of 100%, spatial frequency of 0.002-0.256 cycles/degree, and a temporal frequency of 1-3 Hz. Gratings were presented by stepping the orientation from 0-360 degrees in steps of 30 degrees, with each grating presentation being preceded for 4 seconds “off” followed by 2 seconds “on’.

For quantitative analyses, the OSI was computed by taking the vector average of responses to all orientations, according to the formula described previously (Banerjee et al., 2016):

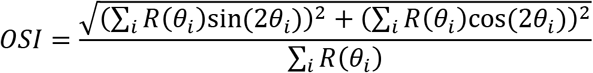

Cells were filtered based on visual responsiveness (t-test, OFF vs. ON, p<0.05) and selectivity (OSI >0.15).

### Slice Electrophysiology

Mice were anesthetized with isoflurane and the brain was rapidly removed and sliced coronally using a vibratome (VT1200S; Leica) at a thickness of 300μm in slicing buffer (mM: 130 choline chloride, 25 glucose, 1.25 NaH2PO4, 26 NaCHO3, 2.5 KCl, 7 MgCl2, and 0.5 CaCl2) bubbled with 95% O2 and 5% CO2. Slices were incubated for a minimum of 60 min in room-temperature ACSF (mM: 130 NaCl, 10 glucose, 1.25 NaH2PO4, 24 NaCHO3, 3.5 KCl, 2.5 CaCl2, and 1.5 MgCl2). For recording of AMPA receptor (AMPAR)-mediated mEPSCs, whole-cell patch clamp of layer II/III pyramidal neurons in the binocular region of V1 was performed using pipettes (4–7 MΩ resistance) filled with an internal solution (mM: 100 K-gluconate, 20 KCl, 0.5 EGTA, 10 NaCl, 10 Na-phosphocreatine, 4 Mg-ATP, 0.3 Na-GTP, and 10 HEPES, pH 7.2–7.3 titrated with 1M KOH). Neurons were recorded at room temperature (25°C) in ACSF containing 1 μM TTX, 50 μM AP-5, and 50 μM picrotoxin to isolate AMPAR-mediated currents and voltage clamped at a membrane potential of −70 mV. mEPSCs were recorded using a Multiclamp 700B amplifier (Molecular Devices) at 10 kHz, filtered at 2 kHz, and analyzed with Clampfit 10.2 software (Molecular Devices). Whole-cell membrane currents were recorded for 10 min. For detection of mEPSCs, a detection template for each cell was constructed from four to six events intrinsic to each recording. Traces were analyzed in template search mode in Clampfit 10.2, with a template match threshold of 4–4.5 to reduce false positives. All events were detected automatically and edited after detection by eye to remove events that were erroneous matches or duplicate events. All mEPSC events were included in the analysis of event parameters.

### Intrinsic Signal Optical Imaging

Mice were anesthetized with urethane (1.5 mg/g, i.p.) and chloroprothixene (10 mg/kg, i.p.). The skull was exposed over V1 and a head plate fixed to the head and minimize movements. The cortex was covered with agarose solution (1.5%) and a glass coverslip. Red light (630 nm) was used to illuminate the cortical surface, and the change of luminance was captured by a CCD camera (Cascade 512B; Roper Scientific) during the presentation of visual stimuli (custom MATLAB scripts). An elongated horizontal white bar (9° × 72°) over a uniformly gray background was drifted upward continuously through the peripheral-central dimension of the visual field. After moving to the last position, the bar would jump back to the initial position and start another cycle of movement; thus, the chosen region of visual space (72° × 72°) was stimulated in a periodic manner (12 s / cycle). Images of the visual cortex were continuously captured at the rate of 18 frames/s during each stimulus session of 22 min. A temporal high-pass filter (135 frames) was used to remove slow noise components, after which the temporal fast Fourier transform (FFT) component at the stimulus frequency (9 s^−1^) was calculated pixel by pixel from the entire set of images. The amplitude of the FFT component was used to measure the strength of visually driven response for each eye, and the OD index (ODI) was derived from the response of each eye (R) at each pixel as ODI = (R_contra_ − R_ipsi_) / (R_contra_+R_ipsi_). The binocular zone was defined as the cortical region that was driven by stimulation of both the ipsilateral and contralateral eyes. The response amplitude for each eye was defined as fractional changes in reflectance over baseline reflectance (ΔR/R ×10^−3^), and the top 50% pixels were analyzed to avoid background contamination.

### Statistical Analyses

Two-tailed unpaired Student’s t-tests were used for comparisons between two means. For comparing more than two means, a one-way or two-way analysis of variance (ANOVA) was used as appropriate followed by post-hoc pairwise comparisons using the Holm-Sidak method. Individual data points plotted represent averages from separate animals. All statistical tests were performed using Prism (GraphPad, La Jolla). All averaged data are presented as mean ± SEM.

## RESULTS

### GLT1 expression in the visual cortex during development

We first confirmed that GLT1 is expressed in the visual cortex during the critical period for ocular dominance plasticity (~P28) when activity driven refinement is particularly prominent. Indeed, we found that GLT1 protein was highly expressed across all cortical layers (**Figure 1A,B**) as previously reported (Massie et al., 2003; Voutsinos-Porche et al., 2003). Next, we asked whether GLT1 transcription changed during a period of eye-specific refinement and binocular matching in V1. GLT1 mRNA transcripts were collected from visual cortex from P0-P60 and normalized to P28 to determine whether GLT1 transcription correlated with eye-opening and synaptic refinement (**Figure 1C**). We found that GLT1 mRNA was developmentally regulated (one-way ANOVA, F(6,18) = 86.7, p=2.7×10^−12^) and that it remained low until ~P14, around the time of eye-opening (P28: 1.0±0.01, P0: 0.03±0.01, P7:0.18±0.01, P14: 0.89±0.02, P21: 1.28±0.14, P42: 1.02±0.08, P60: 1.04±0.03). GLT1 transcripts peaked around P21 and remained high through P60, indicating that GLT1 function could be important for experience-dependent binocular refinement that occurs during the critical period of ocular dominance plasticity between ~P14-34. Furthermore these findings suggest that GLT1 expression may be regulated by visual experience, and thus have a role in driving activity-dependent plasticity in V1.

**Figure 1:**
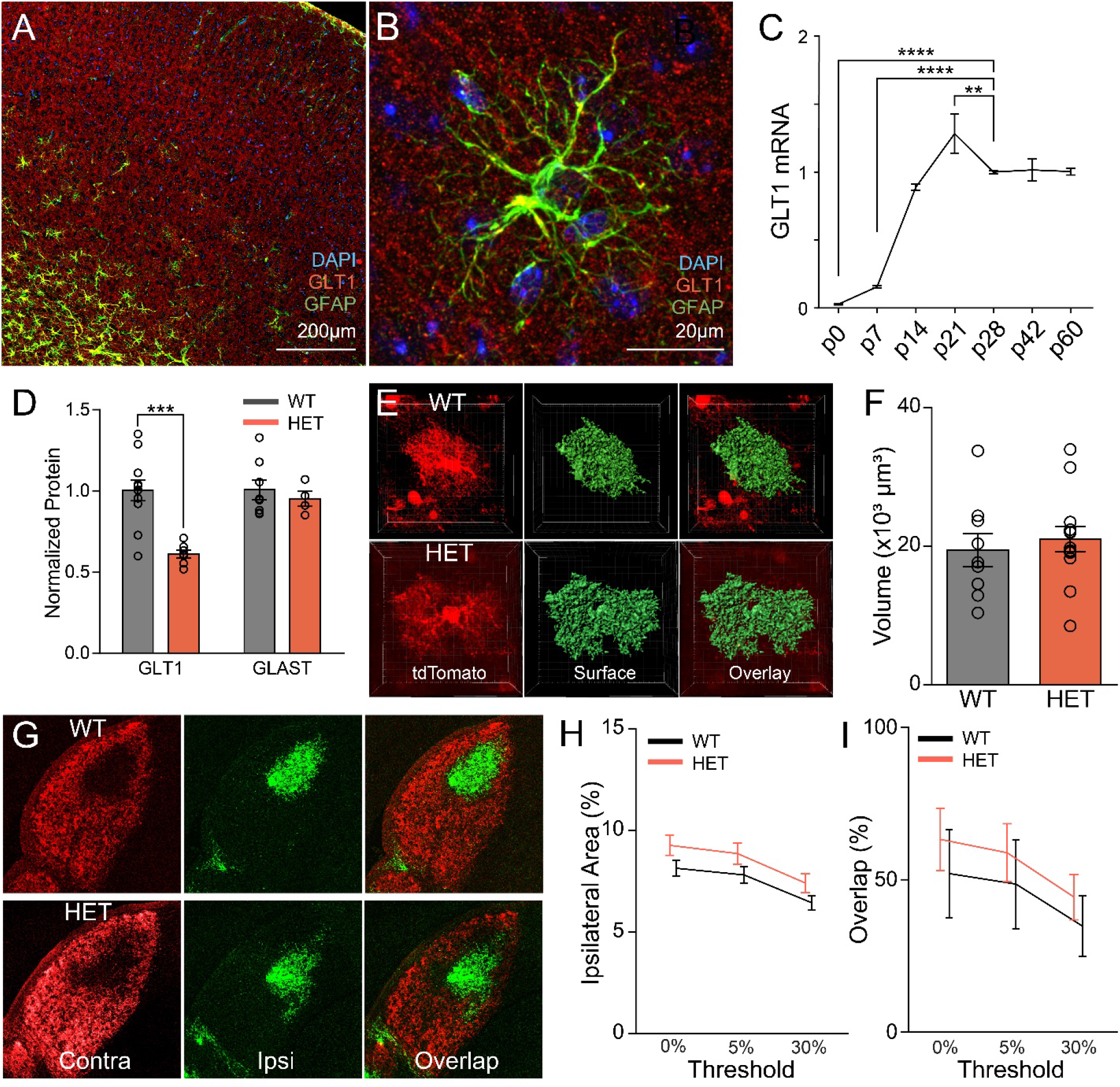
GLT1 is upregulated in the developing visual cortex concurrent with visual experience. **A)** Immunohistochemical stain of GLT1 (red), astrocytes (green), and DAPI (blue) at ~P28 in mouse visual cortex. GLT1 is expressed by astrocytes throughout cortical layers. **B)** High-magnification image of a single GFAP-labelled astrocyte with surrounding GLT1 expression. **C)** Quantification of GLT1-mRNA across developmental time points showing a significant increase from birth and peaking at P21. Levels are normalized to P28 (P0 (n=4), P7 (n=4), P14 (n=3), P21 (n=3), P28 (n=4), P42 (n=3), P60 (n=4); one-way ANOVA, F(6,18)=86.7, p=2.7×10^−12^). **D)** Western blot quantification showing that transgenic mice with heterozygous expression of GLT1 (HET) have significantly less GLT1 expression compared to wildtype (WT) littermates (n=WT(11), HET(11); unpaired t-test, t=4.64, p=2.7×10^−4^), but comparable expression of GLAST (WT (n=8), HET (n=4); unpaired t-test, t=0.62, p=0.55). **E)** Example images of GLT1 WT and HET astrocytes labeled using a custom GFAP-tdTomato transgenic mouse line (red). Astrocyte volume is reconstruction from imaged z-stacks (green). **F)** Quantification of astrocyte volume from 3D reconstructions show no difference between GLT1 WT and HET animals (WT (n=3), HET (n=13); unpaired t-test, t=0.53, p=0.60). **G)** Images of the lateral geniculate nucleus (LGN) after CTB-594 (red) and CTB-488 (green) injection into the contralateral and ipsilateral eyes respectively. Merged overlays from GLT1 WT and HET mice show normal retinothalamic axon segregation. **H)** Quantification of ipsilateral area in GLT1 WT and HETs across several binary thresholds (0, 5, 30%) showing no difference in absolute ipsilateral area (WT (n=3), HET (n=6); two-way ANOVA, genotype effect F(1,7)=1.93, p=0.21). **I)** Quantification of contra/ipsi projection overlap showing no difference in contra/ipsi segregation (WT (n=3), HET (n=6); two-way ANOVA, genotype effect, F(1,7)=0.43, p=0.53). *p<0.05, **p<0.01, ***p<0.005

### GLT1 heterozygous mice have normal astrocyte morphology and retinogeniculate axonal segregation

In order to gain a deeper understanding of the role of GLT1 in visual cortex development, we used a germline heterozygous knockout GLT1 mouse model. This mouse line (GLT1 HET) had been previously described as a model for glutamatergic hyperactivity, with GLT1 HET mice displaying several behavioral phenotypes (Tanaka et al., 1997; Kiryk et al., 2008) but had not been widely used for developmental studies. As expected, GLT1 HET animals expressed significantly reduced levels of GLT1 protein in V1 (WT: 1.0±0.07, HET: 0.61±0.02; unpaired t-test, t=4.64, p=2.7×10^−4^; **Figure 1D**). The other major glutamate transporter expressed in the visual cortex of mice, GLAST, was not altered in the GLT1 HETs indicating that there is no compensatory increase in total transporter protein expression in V1 (WT: 1.0±0.06, HET: 0.95±0.05; unpaired t-test, t=0.62, p=0.55). GLT1 expression has been linked to astrocyte process ramification and synaptic ensheathment (Zhou and Sutherland, 2004; Genoud et al., 2006; Benediktsson et al., 2012), suggesting that a constitutive decrease in GLT1 expression might alter astrocyte morphology. To test this hypothesis, we crossed GLT1 HET mice with a GFAP-tDTomato reporter mouse in which astrocytes express tDTomato throughout the cortex. Confocal stacks of tDTomato+ astrocytes were collected from fixed tissue of GLT1 WT and HET mice and 3D-reconstructed using the Imaris software package (**Figure 1E**). No significant difference was observed in astrocyte volume between GLT1 WT and HET animals (WT: 19.4×10^3^±2.4×10^3^, HET: 21.0×10^3^±1.8×10^3^; unpaired t-test, t=0.53, p=0.60) showing that ~40% reduction in GLT1 expression does not significantly alter astrocyte morphology (**Figure 1F**). As a prelude to examining ocular dominance plasticity in V1, we also investigated whether GLT1 HET mice showed deficits in the activity-dependent segregation of retinogeniculate axons in the lateral geniculate nucleus (LGN). Cholera toxin subunit B conjugated to two distinct fluorophores was individually injected into the contralateral and ipsilateral eyes labeling the eye-specific projections to the LGN (**Figure 1G**). We found no significant difference between either the ipsilateral axonal area normalized to the contralateral axonal area (two-way ANOVA, genotype effect, F(1,7)=1.93, p=0.21; **Figure 1H**) nor the overlap between the contralateral and ipsilateral axonal areas (two-way ANOVA, genotype effect, F(1,7)=0.43, p=0.53; **Figure 1I**). These results demonstrate that GLT1 HET mice have reduced GLT1 expression in V1, but do not exhibit gross morphological deficits in astrocytes or axonal segregation of retinogeniculate projections in the LGN, and hence in the anatomical extent of retinal drive to visual cortex.

### Visual cortex neurons in GLT1 HET mice have abnormally high ipsilateral responses and poor binocular matching of preferred orientation

We next examined the functional consequences of GLT1 reduction on experience-dependent refinement of binocular inputs to the visual cortex. During normal development, binocular cells in the contralateral cortex that respond to visual input from both the contralateral and ipsilateral eyes (**Figure 2A**) refine their activity to (a) scale responses from the two eyes, and (b) match their preferred activity from both eyes to similarly specific orientations (Gu and Cang, 2016; Hooks and Chen, 2020). To test whether GLT1 HET mice have alterations in the magnitude and refinement of binocular inputs, we examined several measures of neuronal responses to visual gratings stimulating either the contralateral or ipsilateral eye (**Figure 2A**). We used two-photon microscopy to image neuronal responses in V1 of GLT1 WT or HET mice at ~P28. We stimulated the contralateral and ipsilateral eyes independently with gratings of different orientations and examined the calcium responses (**Figure 2B**). We then calculated independent tuning curves to grating orientation for both contralateral and ipsilateral input. We qualitatively observed striking differences between GLT1 WT and GLT1 HET mice. Neuronal responses to gratings in GLT1 WT mice had similar tuning curves to gratings presented to either the contralateral or ipsilateral eye, showing the expected matching of preferred orientation between the two eyes. In addition, responses to contralateral input was higher than ipsilateral input as expected for a normal contralateral bias in responses (**Figure 2B**, top). Conversely, neuronal responses to gratings in GLT1 HET mice had qualitatively mismatched tuning between contralateral and ipsilateral eye input, and abnormally high ipsilateral response magnitudes (**Figure 2B**, bottom). To quantify the neuronal responses between genotypes, we first examined whether there was a difference in the magnitude of the response to the preferred orientation (**Figure 2C**) regardless of the specific preferred orientation angle. We found that, as expected, the average response to preferred orientation of binocular neurons was significantly higher to contralateral inputs relative to ipsilateral inputs in GLT1 WT mice (WT-C/I: 0.30±0.02 / 0.22±0.01, HET-C/I: 0.32±0.03 / 0.33±0.03; one-way ANOVA, F(3,674)=6.44, p=2.7×10^−4^), demonstrating the contralateral bias (Holm-Sidak, t=3.39, p=4.0×10^−3^). However, GLT1 HET neurons displayed comparable response magnitudes to contralateral and ipsilateral input (Holm-Sidak, t=0.25, p=0.85). These response magnitudes were significantly higher than ipsilateral response magnitudes (Holm-Sidak, t=3.42, p=3.9×10^−3^) and comparable to contralateral response magnitudes in GLT1 WT mice (Holm-Sidak, t=0.80, p=0.81; Figure 2C). To directly compare the ratio of responses to contralateral and ipsilateral stimuli, we calculated the ocular dominance index (ODI) for each neuron across genotypes (see methods for ODI calculation, **Figure 2D**). We found that neurons in GLT1 HET mice had a significantly decreased ODI indicating a decrease in the normal contralateral bias (WT: 0.14±0.02, HET: 0.01±0.04; unpaired t-test, t=3.15, p=1.8×10^−3^).

**Figure 2.**
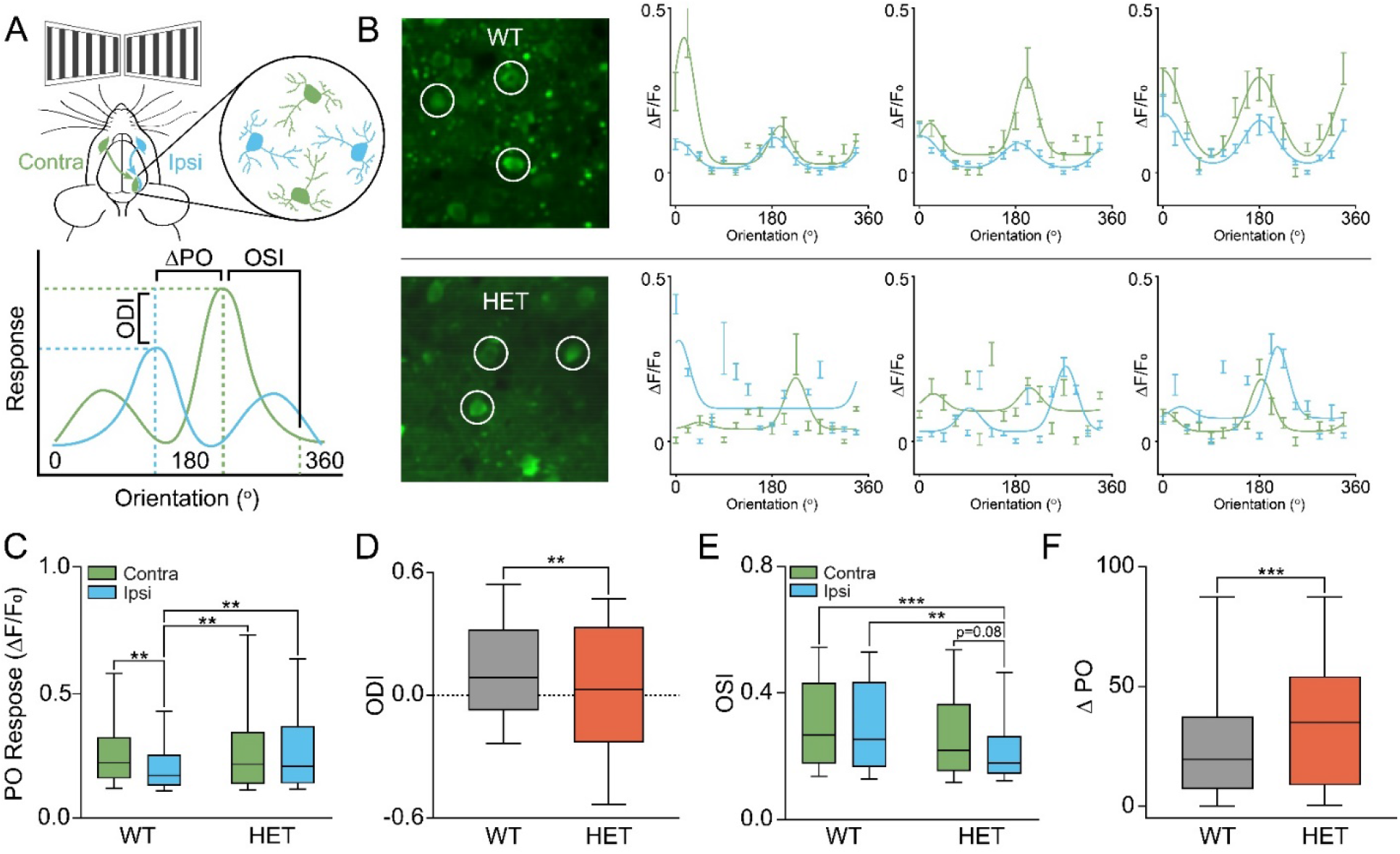
GLT1 HET mice have higher ipsilateral eye responses, lower contralateral eye bias and disrupted experience-dependent binocular matching of orientation-selective responses. **A)** Schematic of experimental design. Top: Visual gratings were separately presented to the contra (green) and ipsi (blue) eyes in P28 mice and neuronal responses recorded. Bottom: schematic of measures. Ocular dominance index (ODI) was calculated as (max^Contra^ − max^Ipsi^) / max^Contra^+max^Ipsi^. Orientation Selectivity Index (OSI) was calculated as described previously (Banerjee et al., 2016). Difference in preferred orientation (ΔPO) was calculated as the difference between preferred orientations of the max contralateral and ipsilateral responses. **B)** Example cells in GLT1 WT (top) and GLT1 HET (bottom) animals. Left: *in vivo* images of neuronal somas measured in binocular visual cortex using the calcium indicator, GCaMP6f. Right: Tuning curves of three cells (white circles in left) to contra (green) and ipsi (blue) stimulation. Note the matched tuning and contralateral bias in WT animals and the mismatched tuning curves and lack of contralateral bias in GLT1 HETs. **C)** Quantification of the average response to PO in GLT1 WT and HET mice. WT mice have a significantly higher contralateral response than ipsilateral response while HET mice have approximately equal contralateral and ipsilateral responses (n=4-6 animals, 23-52 cells per animal, two-way ANOVA, genotype F(1,674)=7.72, p=0.0056, interaction F(1,674)=4.243, p=0.040). **D)** Quantification of ocular dominance index showing that GLT1 HET mice have significantly decreased ODI (n=4-6 animals, 23-52 cells per animal, t-test, p=0.0018). **E)** Quantification of OSI showing that GLT1 HET mice have a significantly decreased OSI of ipsilateral responses compared to both contra and ipsi responses in GLT1 WT animals (n=4-6 animals, 23-52 cells per animal, two-way ANOVA, genotype F(1,674)=12.46, p=4.5×10^−4^). **F)** Quantification of ΔPO showing an increased difference in the preferred orientations between contralateral and ipsilateral inputs to neurons in GLT1 HET animals (n=4-6 animals, 23-52 cells per animal, t-test, p=1.0×10^−4^). *p<0.05, **p<0.01, ***p<0.005

We next asked whether the orientation specificity to tuned responses was different between GLT1 WT and HET mice. We calculated the global orientation selectivity index (OSI) according to previous reports (Banerjee et al., 2016) for each neuron and for contralateral and ipsilateral visual stimulation (**Figure 2E**). We found that in GLT1 WT mice, the OSI was comparable for both contralateral and ipsilateral eye responses (WT-C: 0.31±0.01, WT-I: 0.30±0.01; one-way ANOVA, F(3,674)=6.12,p=4.1×10^−4^; Holm-Sidak, t=0.78, p=0.65). However, in GLT1 HET mice, ipsilateral responses had a significantly lower OSI compared to both contralateral (HET-I: 0.23±0.01, WT-C: 0.31±0.01; Holm-Sidak, t=4.16, p=1.1×10^−4^) and ipsilateral responses (HET-I: 0.23±0.01, WT-I: 0.30±0.01; Holm-Sidak, t=3.56, p=2.0×10^−3^) in WT mice, and a trend towards decreased OSI relative to contralateral responses in GLT1 HET mice (HET-C: 0.28±0.02, HET-I: 0.23±0.01; Holm-Sidak, t=2.30, p=0.08). These results demonstrate that ipsilateral responses in GLT1 HET mice are less specific to orientation tuning than WT mice. Finally, to quantify the degree to which contralateral and ipsilateral peak responses were tuned to the same orientation, indicating binocular matching, we calculated the difference in preferred orientation angle between peak contralateral and ipsilateral input (ΔPO; **Figure 2F**). We found that GLT1 HET neurons had significantly higher difference in preferred orientation angle compared to GLT1 WT neurons (WT: 24.3±1.37, HET: 35.1±2.71; unpaired t-test, t=3.93, p=1.0×10^−4^). Together, these results reveal that in GLT1 HET mice, ipsilateral eye responses are abnormally high leading to reduced ODI, decreased OSI, and an increased in ΔPO, altogether signifying increased ipsilateral drive and disrupted binocular matching.

### GLT1 HET mice have abnormal excitation and inhibition in V1

Given the observed changes in response magnitude and specificity in GLT1 HET mice, we next asked whether changes in structural features of layer 2/3 V1 neurons and in synaptic transmission might explain the neurophysiological changes. We reasoned that if eye-specific responses and binocular matching were disrupted by excess extracellular glutamate, synapses that were normally pruned in an activity-dependent manner may be abnormally retained. To examine this possibility, we crossed the GLT1 HET mice with a Thy1-GFPm mouse line (Feng et al., 2000) that expresses GFP in a subset of neurons across the brain. We chose this line because labeling of layer 2/3 neurons in the visual cortex is relatively sparse, which allowed us to examine synaptic spine density on clearly labeled dendrites (**Figure 3A**). We collected brain slices from ~P28 GLT1 WT and HET mice and analyzed the dendritic spine density on basal dendrites of layer 2/3 excitatory neurons (**Figure 3B**). In addition to qualitative differences in dendrite morphology, we found a significant increase in dendritic spine density (WT (n=4): 0.92±0.09, HET (n=4): 1.18±0.04; unpaired t-test, t=2.60, p=0.04) (**Figure 3C**). We next asked whether the increased dendritic spine density resulted in an increase in functional synapses as measured by spontaneous excitatory post-synaptic currents (mEPSCs) in V1 neurons in slices from GLT1 WT and HET mice at ~P28. Layer 2/3 neurons were whole-cell patched and mEPSCs measured (**Figure 3D**). We found no difference in the amplitude of mEPSCs between GLT1 WT and HET neurons (WT (n=13 cells): −11.1±0.38, HET (n=8 cells): −11.6±0.49; unpaired t-test, t=0.70, p=0.49; **Figure 3E**), however there was a trend toward increased frequency suggesting that GLT1 HET mice had increased excitatory input (WT (n=13 cells): 0.92±0.08, HET (n=8 cells): 1.28±0.18; unpaired t-test, t=2.07, p=0.05; **Figure 3F**).

**Figure 3.**
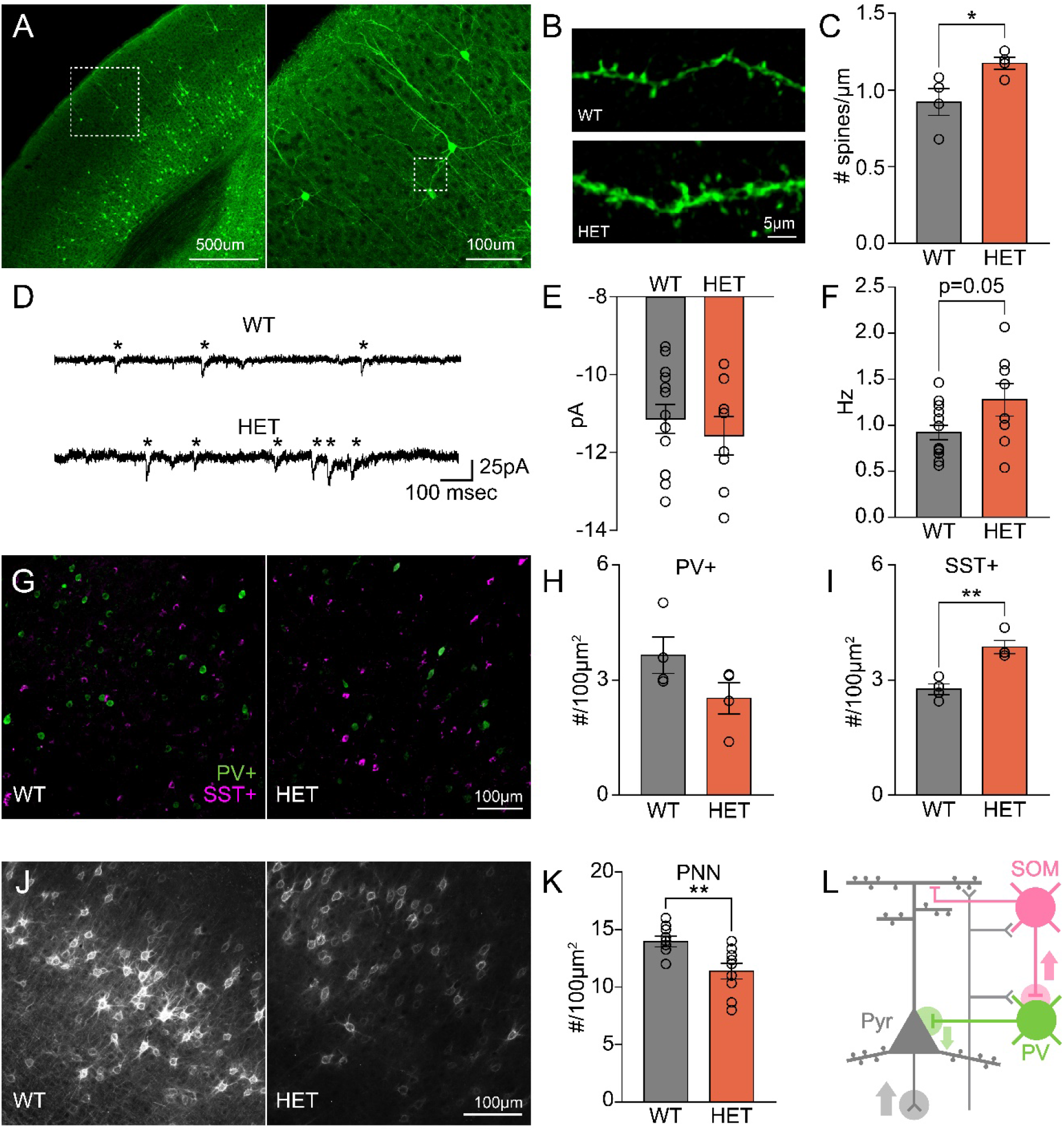
GLT1 HET mice have abnormal maturation of excitatory and inhibitory circuits. **A)** Left: low-magnification image of neurons in visual cortex of GFP-M transgenic mice. Right: higher-magnification image of layer 2/3 neurons (dotted box in left). **B)** Images of basal dendrites of layer 2/3 neurons in GLT1 WT (top) and HET (bottom) mice. WT example is from dotted box in right panel of A. **C)** GLT1 HET mice have increased spine density on basal dendrites of layer 2/3 neurons in visual cortex (n=4 animals, 5 slices, 10 dendrites per animal, t-test, p=0.041). **D)** Example traces of miniature excitatory post-synaptic currents (mEPSCs) from voltage-clamped layer 2/3 neurons in the visual cortex of GLT1 WT and HET mice. **E)** Quantification of mEPSC amplitude showing no difference in magnitude of mEPSCs (n=8-13 cell, t-test, p=0.49). **F)** Neurons from GLT1 HET mice have a trend towards increased mEPSC frequency (n=8-13 cells, t-test, p=0.052). **G)** Example images of parvalbumin positive (PV+, green) and somatostatin positive (SST+, magenta) interneurons in visual cortex of GLT1 WT and HET mice). **H)** GLT1 HET mice have a trend towards decreased PV+ neuron density (n=4 animals, 5 slices per animal, t-test, p=0.12). I) GLT1 HET mice have a significant increase in SST+ cell density (n=4 animals, 5 slices per animal, t-test, p=0.0023). **J)** Example images of perineuronal nets (PNNs) visualized using wisteria floribunda agglutin (WFA) staining. **K)** GLT1 HET mice have significantly decreased PNN density compared to WT littermates (n=9 animals, 5 slices per animal, t-test, p=0.0068). **L)** Model of net decrease in cortical inhibition through increased SST+ cell density inhibiting PV+ interneurons yielding increase in excitatory pyramidal neuron responses (Pyr). *p<0.05, **p<0.01,***p<0.005

Increased dendritic spine density and excitatory drive may enhance excitation and also alter the balance of excitation and inhibition in cortical circuits. Specifically, we reasoned that changes in glutamate uptake may affect excitatory neurotransmission onto inhibitory-disinhibitory neuron classes. In the visual cortex, excitation is balanced with inhibition mediated by specific interneuron classes that selectively target pyramidal neuron somata and dendrites, as well as other inhibitory neurons (Kepecs and Fishell, 2014). The number and density of inhibitory neurons is remarkably sensitive to changes in input drive (Lazarus and Huang, 2011; Hooks and Chen, 2020). We thus examined whether GLT1 HET mice had alterations in the density of two main classes of interneurons, parvalbumin positive (PV) and somatostatin positive (SST) neurons. We stained coronal slices of primary visual cortex from GLT1 WT and HET mice for PV and SST using dual-color immunohistochemistry (**Figure 3G**). We then examined PV or SST cell density in the binocular region (according to anatomical estimates) of V1 and found that although PV cell density was not significantly different (WT (n=4): 6.09±0.79, HET (n=4): 4.22±0.68; unpaired t-test, t=1.78, p=0.12; **Figure 3H**), SST neuron density was significantly higher in GLT1 HET mice compared to WT mice (WT (n=4): 2.76±0.14, HET (n=4): 3.86±0.17; unpaired t-test, t=5.05, p=2.3×10^−3^; **Figure 3I**). The maturation of PV neurons and their inhibitory input onto excitatory somas of pyramidal neurons are hallmarks of experience-dependent plasticity and synaptic refinement (Yazaki-Sugiyama et al., 2009; Hooks and Chen, 2020). Specifically, the formation of perineuronal nets (PNNs) around pyramidal somas is thought to reflect the closing of the critical period (Katagiri et al., 2007). Given the abnormal density of SST neurons, which are known to inhibit PV neurons, we examined whether there was a difference in PNN density indicating shifts in inhibitory maturation. We used wisteria floribunda agglutinin (WFA) to stain PNNs in binocular visual cortex of GLT1 WT and HET mice (**Figure 3J**). Quantification of PNN density revealed a significant decrease in GLT1 HET mice compared to WT controls suggesting a decreased maturation of somatic inhibition WT (n=9): 13.95±0.46, HET (n=9): 11.38±0.69; unpaired t-test, t=3.11, p=6.8×10^−3^; **Figure 3K**). Overall, these results suggest that GLT1 HET mice have increased excitatory synaptic drive to layer 2/3 neurons, along with decreased somatic inhibition on pyramidal neurons (**Figure 3L**), which together may help mediate the altered magnitude and tuning of their eye-specific responses.

### GLT1 HET mice have disrupted ocular dominance plasticity

Our observations suggest that experience-dependent synaptic refinement underlying ipsilateral response amplitude and binocular orientation matching is altered in GLT1 HET mice. Therefore, we asked whether experience-dependent synaptic plasticity is broadly altered in GLT1 HET mice. To address this question, we used the classic ocular dominance plasticity (ODP) model of synaptic plasticity in which one eye is deprived of visual input for 4-7 days during the critical period for ODP (~P21-P35 in mice) resulting in a broad network shift in the contralateral visual cortex of preferred responsiveness to the deprived eye (typically the contralateral eye) toward the eye that remained open (typically the ipsilateral eye) (Hooks and Chen, 2020). Previous work has demonstrated that 7 days of monocular deprivation (MD) leads first to a decrease in responsiveness to the closed eye in the first 3-4 days followed by potentiation of responsiveness to the open eye (Smith et al., 2009). The degree to which this shift occurs indicates the degree to which cortical circuits are able to functionally and structurally rearrange in response to changes in sensory input. To assess ODP in GLT1 WT and HET mice, we used intrinsic signal optical imaging, which has been used to characterize large-scale neuronal response to visual stimuli across V1 (Cang et al., 2005). GLT1 WT and HET mice at ~P28 were either not deprived (ND), or monocularly deprived for 4 days (4dMD) or 7 days (7dMD). Contralateral and ipsilateral eyes were then stimulated individually, and V1 contralateral to the deprived eye imaged. A drifting bar was used to evoke a retinotopic map of activity and the binocular region was selected to evaluate the amplitude of response elicited by contralateral and ipsilateral eye stimulation (**Figure 4A**).

**Figure 4.**
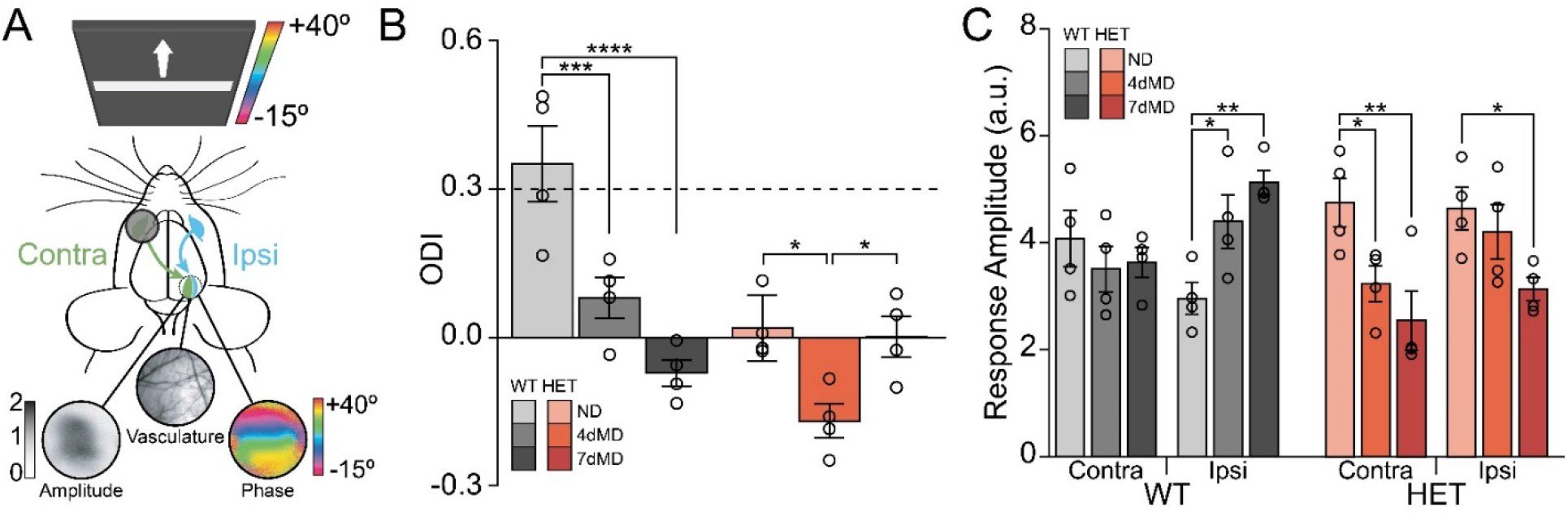
GLT1 HET mice have disrupted ocular dominance plasticity. **A)** Schematic of experimental setup for intrinsic signal optical imaging. Drifting bars are presented to each eye individually and phase maps are generated by the retinotopic activity in visual cortex. Averaged responses of multiple sweeps yield an amplitude map. Ocular Dominance Index (ODI) is calculated as the contralateral response − ipsilateral response / contralateral + ipsilateral responses. **B)** ODI for GLT1 WT (grays) and GLT1 HET (reds) mice that were either non-deprived (ND), or had the contralateral eye monocularly deprived for 4 days (4dMD) or 7 days (7dMD). GLT1 WT mice display a typical contralateral bias in ND conditions. After 4dMD and 7dMD the ODI significantly decreases demonstrating intact ocular dominance plasticity. GLT1 HET mice display an abnormal lack of contralateral ODI bias under ND conditions, a significant decrease in ODI at 4dMD, and a return to no bias at 7dMD (n=4 animals per group, two-way ANOVA, Holm-Sidak post-hoc comparisons). **C)** Comparison of eye-specific amplitudes for GLT1 WT and HET mice. ND GLT1 HET mice have approximately equal responses to contralateral and ipsilateral inputs. After 4dMD, GLT1 HET mice have a significant decrease in contralateral responses and at 7dMD, significant decrease in both contralateral and ipsilateral responses (n=4 animals per group, two-way ANOVA, Holm-Sidak post-hoc comparisons). *p<0.05, **p<0.01, ***p<0.005

We calculated the ODI of imaged pixels to evaluate relative responses to contralateral and ipsilateral eye stimulation. We first recapitulated normal ODP in ND GLT1 WT mice as evidenced by a strong contralateral bias (ODI>0.3, **Figure 4B**). Following 4dMD, GLT1 WT mice displayed a normal ocular dominance shift with significant reduction in ODI (WT-ND (n=4): 0.35±0.04, WT-4dMD (n=4): 0.05±0.02; Holm-Sidak, t(18)=4.83, p=2.7×10^−4^). Following 7dMD we observed a further significant shift in ODI resulting in a slight ipsilateral bias (WT-ND (n=4): 0.35±0.04, WT-7dMD (n=4): −0.07±0.01; Holm-Sidak, t(18)=6.83, p=6.5×10^−6^). Next, we repeated the imaging in GLT1 HET mice and found that unlike WT controls, ND HET animals showed a significantly reduced ODI consistent with our single-cell calcium recordings (see Figure 2D). Following 4dMD, GLT1 HET mice showed a strong ocular dominance shift resulting in a pronounced ipsilateral bias, and demonstrating that ODP was intact (HET-ND (n=4): 0.02±0.02, HET-4dMD (n=4): −0.17±0.02; Holm-Sidak, t(18)=3.03, p=0.02). Surprisingly, after 7dMD GLT1 HET mice had a significant ocular dominance shift back towards contralateral responsiveness, indicating plasticity opposite to the direction of experience-dependent potentiation (HET-4dMD (n=4): −0.17±0.02, HET-7dMD (n=4): 0.002±0.02; Holm-Sidak, t(18)=2.76, p=0.03). In order to explain these counterintuitive results, we examined the individual response amplitudes to each eye across 4dMD and 7dMD. In GLT1 WT mice, we observed a reduction in responses to the contralateral eye and a significant increase in responsiveness to the ipsilateral eye consistent with previous work (WT-ND-I (n=4): 3.01±0.15, WT-4dMD-I (n=4): 4.4±0.24; Holm-Sidak, t(36)=2.48, p=0.04; WT-ND-I (n=4): 3.01±0.15, WT-7dMD-I (n=4): 5.13±0.11; Holm-Sidak, t(36)=3.74, p=1.9×10^−3^)(Sato and Stryker, 2008). However, in GLT1 HET mice, we observed an overall decrease in contralateral and ipsilateral responsiveness at both 4dMD and 7dMD with the contralateral response decreasing in amplitude at 4dMD (HET-ND-C (n=4): 4.75±0.23, HET-4dMD-C: 3.24±0.17; Holm-Sidak, t(36)=2.63, p=0.02) and 7dMD (HET-ND-C (n=4): 4.75±0.23, HET-7dMD-C: 2.55±0.28; Holm-Sidak, t(36)=3.81, p=1.6×10^−3^), and ipsilateral response at 7dMD (HET-ND-I (n=4): 4.65±0.20, HET-7dMD-I: 3.14±0.11; Holm-Sidak, t(36)=2.60, p=0.04). These results suggest that GLT1 HET mice show a normal decrease in contralateral responsiveness, but an abnormal decrease in ipsilateral responsiveness with MD as well. Therefore, the increase in ODI from 4dMD to 7dMD in GLT1 HETs reflects a predominating decrease in ipsilateral responses rather than an increase in contralateral responses.

### GLT1 expression in GLT1 HET mice is modulated by monocular deprivation

Previous work has demonstrated that GLT1 protein expression and translocation are influenced by changes in neuronal activity (Benediktsson et al., 2012; Murphy-Royal et al., 2015). Although GLT1 HET mice have ~40% reduction in protein at ~P28 in the absence of MD, it is possible that MD affects GLT1 expression in the remaining copy of the GLT1 gene. To assess this possibility, we measured GLT1 mRNA levels in GLT1 WT and HET mice either in ND or 4dMD and 7dMD conditions (**Figure 5A**). In ND mice, we observed the expected decrease in GLT1 HET mice relative to WT controls (WT-ND (n=2): 1.00±0.02, HET-ND (n=3): 0.60±0.02; Holm-Sidak, t(9)=2.95, p=0.02). Interestingly, in GLT1 HET mice, we observed a significant increase in GLT1 mRNA at 4dMD (HET-ND (n=3): 0.60±0.04, HET-4dMD (n=3): 1.16±0.04; Holm-Sidak, t(9)=4.60, p=2.6×10^−3^) and 7dMD (HET-ND (n=3): 0.60±0.04, HET-7dMD (n=3): 1.30±0.17; Holm-Sidak, t(9)=4.60, p=2.6×10^−3^), suggesting that MD increases GLT1 expression in a haplosufficient manner. To determine whether increases in GLT1 mRNA lead to increased GLT1 protein, we performed western blot analysis of GLT1 protein in GLT1 WT and HET mice at either ND, 4dMD, or 7dMD (**Figure 5B**). We reproduced our previous data showing an ~40% reduction in GLT1 protein in ND GLT1 HET mice compared to WT controls (WT-ND (n=9): 1.0±0.02, HET-ND (n=7): 0.61±0.01; Holm-Sidak, t(28)=5.42, p=1.3×10^−4^). However, GLT1 HET mice had an increase in GLT1 protein following 4dMD (HET-ND (n=7): 0.61±0.01, HET-4dMD (n=4): 0.91±0.02; Holm-Sidak, t(28)=3.39, p=0.02) and 7dMD (HET-ND (n=7): 0.61±0.01, HET-7dMD (n=4): 0.92±0.05; Holm-Sidak, t(28)=3.74, p=0.01). The fact that GLT1 WT mice did not have a significant increase in GLT1 protein with MD indicates that increased GLT1 in HET mice with MD is a haploinsufficient phenotype. Importantly, these data reveal that GLT1 HET mice specifically have an increase in GLT1 protein to WT levels in response to MD, which may underlie their abnormal ocular dominance plasticity after MD.

**Figure 5.**
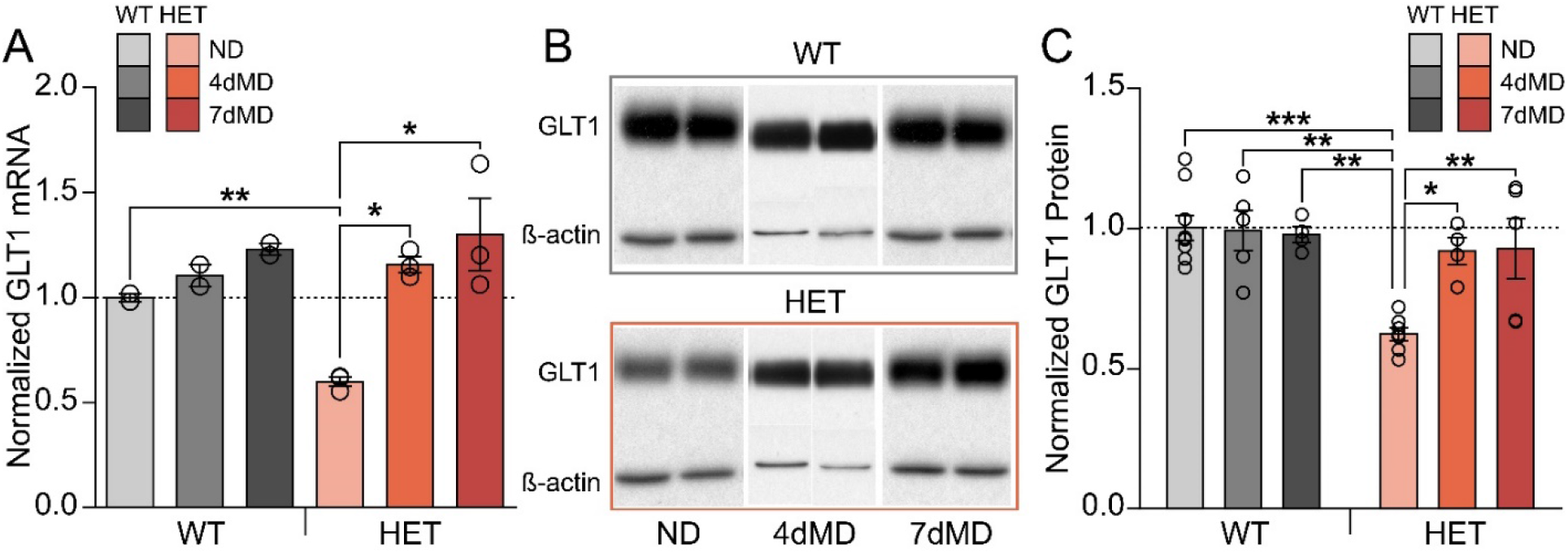
GLT1 HET mice have upregulation of GLT1 expression during monocular deprivation. **A)** Quantification of GLT1 mRNA in WT (grays) and HET (reds) mice in ND, 4dMD, and 7dMD conditions. GLT1 HET mice have significantly less GLT1 mRNA in ND conditions, but no difference at 4dMD and 7dMD compared to WT mice (n=2-3 animals per group, two-way ANOVA, Holm-Sidak post-hoc comparisons). **B)** Example western blots for GLT1 protein in WT (top) and HET (bottom) mice in ND, 4dMD, and 7dMD conditions. **C)** Quantification of western blots showing significantly less GLT1 protein in HET mice in ND, but no difference in 4dMD and 7dMD compared to WT littermates (n=4-9 animals per group, two-way ANOVA, Holm-Sidak post-hoc comparisons). *p<0.05, **p<0.01, ***p<0.005

## DISCUSSION

In summary (**Figure 6**), we confirm that astrocytes in the visual cortex express high levels of GLT1 protein in the visual cortex at ~P28. GLT1 transcription rises in the visual cortex around eye-opening and the onset of experience-dependent synaptic refinement, peaks around P21 and remains high into adulthood. ND GLT1 HET mice have ~40% reduction in GLT1 protein at P28 and normal astrocyte morphology and retinal axonal segregation in the LGN. Single-cell visual responses in the binocular V1 at ~P28 reveal that GLT1 HET mice have abnormally high ipsilateral eye responses, concomitant with increased discrepancy of preferred orientation angle between contralateral and ipsilateral responses, decreased ipsilateral OSI, and decreased ODI. Histological characterization of GLT1 HET mice demonstrates increased dendritic spine density in layer 2/3 pyramidal neurons, increased density of SST+ interneurons, and decreased density of PNNs. Finally, GLT1 HET mice show a robust ocular dominance shift at 4dMD consistent with reduction of deprived, contralateral eye responses. However, they show a paradoxical decrease in nondeprived, ipsilateral eye responses at 7dMD, resulting in an atypical positive shift in ODI. This reduction of ipsilateral eye responses follows an increase in GLT1 protein expression after MD. These findings reveal two surprising features of activity-dependent development in the mouse primary visual cortex that are influenced by astrocyte glutamate transporters: increased ipsilateral eye response magnitude during typical visual experience concurrent with reduced GLT1 expression, and an anomalous effect of MD on eye-specific responses with a reduction in ipsilateral eye responses, concurrent with increased GLT1 expression.

**Figure 6.**
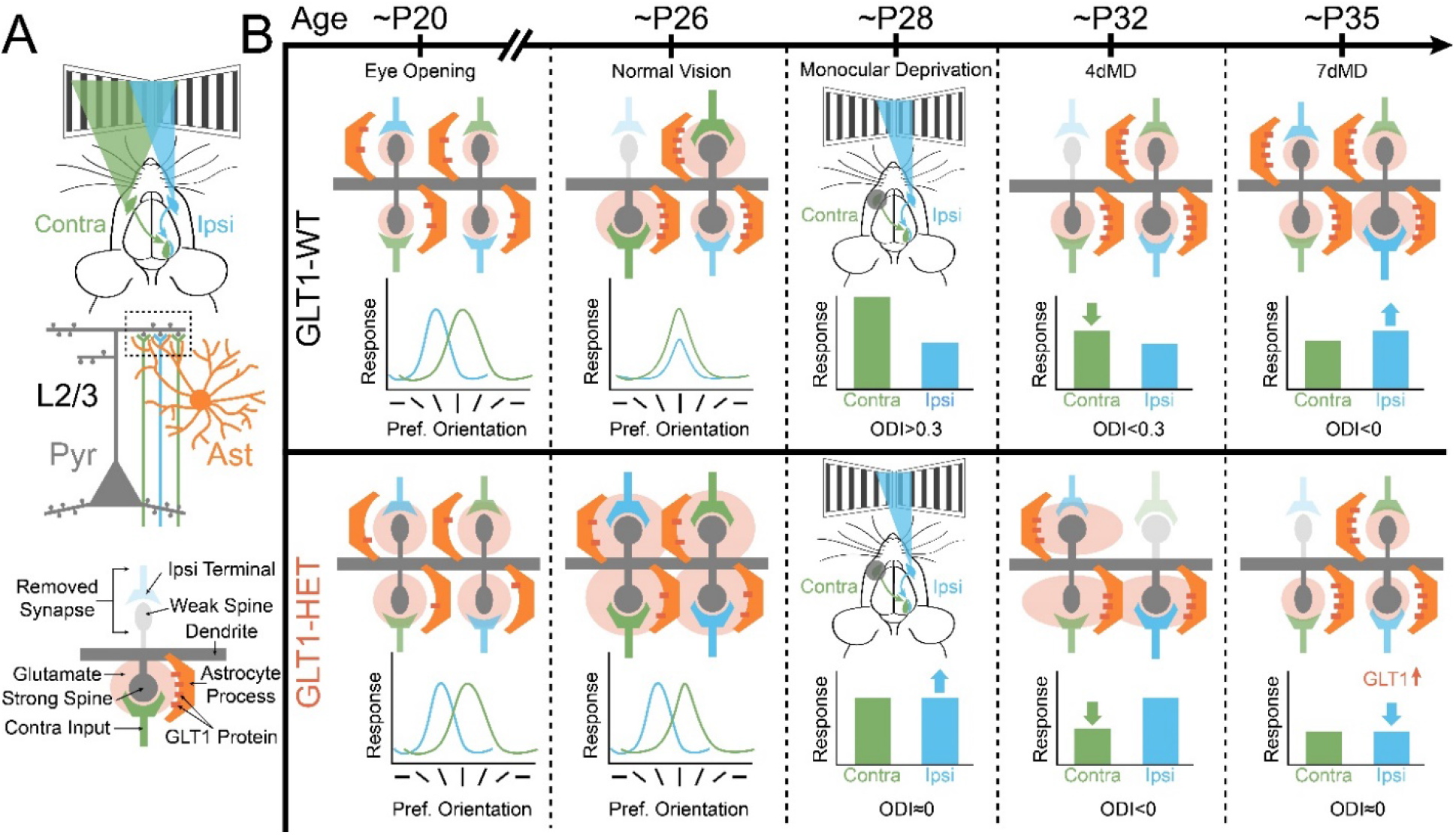
Summary and model of experimental results. **A)** Schematic showing the contralateral (contra, green) and ipsilateral (ipsi, blue) inputs to binocular visual cortex synapsing onto layer 2/3 pyramidal cells (L2/3, Pyr, gray). Astrocytes (Ast, orange) have fine processes that surround excitatory synapses. Dashed box is magnified below, with details. **B)** At eye-opening, GLT1 WT and HET animals have similar astrocyte volume and LGN refinement. With visual experience, GLT1-WT animals undergo activity-dependent plasticity resulting in decreased spine density, binocular matching of preferred orientation, and a contralaterally biased ocular dominance. GLT1-HET animals have comparatively decreased GLT1 protein resulting in increased dendritic spines, increased ipsi responses, reduced contra bias and orientation tuning, and decreased binocular matching of orientation preference. Following 4 days of MD in GLT1-WT mice, contralateral responses decrease and after 7 days of MD, ipsilateral responses increase. In GLT1-HET animals, after 4 days of MD contralateral responses decrease resulting in a negative ODI. However, after 7 days of MD, increased GLT1 expression also decreases ipsilateral inputs resulting in no ocular dominance bias. These results are reasonably explained by a selective influence of GLT1 on ipsilateral inputs and responses during development.

Previous work has demonstrated that non-neuronal glia cells are critical players in both early synaptic refinement and adolescent plasticity including microglial roles in retinothalamic axonal pruning and synaptic remodeling in ocular dominance plasticity (Schafer et al., 2012; Sipe et al., 2016; Stowell et al., 2019). With regards to astrocytic roles in visual cortex development, evidence suggests that astrocytes are able to drive the plasticity of neuronal circuits (Muller and Best, 1989; Foxworthy and Medina, 2015; Singh et al., 2016). Though astrocytes are known to release neuroactive factors to promote synapse formation, our results show that glutamate uptake via GLT1 is another crucial function of astrocytes in synaptic development and plasticity.

The unexpected result that GLT1 reduction leads to abnormally high ipsilateral responses, broader tuning, and mismatched binocular orientation preference can be reasonably explained by: (a) excess glutamate availability at synapses, increased glutamate spillover, and aberrant synaptic pruning; and (b) spatiotemporal control of glutamate at excitatory synapses on features of visual cortex development, including binocular response magnitude and orientation matching. It is possible that distinct cellular mechanisms differentially regulate contralateral and ipsilateral responses and plasticity. For example, it has recently been shown that Homer1a specifically and actively establishes the contralateral bias intrinsic to binocular-responsive cells (Chokshi et al., 2019). Our work indicates that ipsilateral response depression may be more susceptible to changes in GLT1-mediated glutamate uptake. Further work will be needed to explore whether GLT1 expression is differentially regulated at eye-specific synapses.

GLT1 HET mice show a disruption in inhibitory/disinhibitory circuits that may underlie the increase in synaptic spines and aberrant ipsilateral responses. Previous work has established that the maturation of PV+ interneurons, including the chondroitin sulfate proteoglycan PNNs that surround PV+ somas are critical in regulating the closure of the critical period (Pizzorusso et al., 2002; Lensjo et al., 2017; Faini et al., 2018). We find that in GLT1 HET mice, there is a significant decrease in PNNs, suggesting that GLT1 HET mice are uncharacteristically plastic, allowing for increased excitation by aberrant ipsilateral responses. In addition, we find a significant increase in SST+ interneurons, which are known to target PV+ interneurons (Pfeffer et al., 2013) and may aberrantly disinhibit pyramidal neurons. Future studies will be required to determine whether PNNs and SST+ densities eventually return to normal levels later in GLT1 HETs.

Perhaps the most surprising result of our study is the paradoxical observation that MD leads to reduction of non-deprived ipsilateral eye responses following 7dMD. This effect is overlaid on an increased magnitude of ipsilateral eye responses in ND, or pre-MD, conditions. It may be reasonably explained by a homeostatic increase in GLT1 levels in HET animals after MD, and a subsequent specific effect on ipsilateral eye responses that works to re-establish contralateral bias. The fact that ipsilateral deprived eye responses decrease rather than show their usual potentiation after 7dMD supports the idea that GLT1 upregulation is potently coupled to altered synaptic drive, and surprisingly, serves specifically to reduce ipsilateral eye drive relative to contralateral eye drive. It is possible that the switch from genetically driven synaptic organization to experience-dependent synapse refinement marks the upregulation of GLT1 protein in the visual cortex. Although we did not test whether GLT1 reduction affects synaptic plasticity in other sensory cortices, such as the auditory and somatosensory systems, it is possible that GLT1 expression would also occur concomitantly with the activity-dependent regulation of synaptic refinement in those systems.

## ACKNOWLEDGMENTS

We thank Taylor Johns, Vincent Pham, Liadan Maire, Stephanie Chou, and Travis Emery for technical assistance, and Kohichi Tanaka and Jeffrey Rothstein for the GLT1 HET mice. Supported by 1F32EY028028 to GOS, 1F32EY022264 to JP, and NIH grants R01EY028219 and R01DA049005 to MS.

## Notes

### Competing Interest Statement

The authors have declared no competing interest.

